# Protein-protein interaction map in pre-eclampsia through the integration of hub genes, transcription factors and microRNAs

**DOI:** 10.1101/2022.02.28.482425

**Authors:** Praveen Kumar Guttula, Kirti Agrawal, Neelesh Sharma, Mukesh Kumar Gupta

## Abstract

Pre-eclampsia causes complications in pregnancy and characterized by uremia, proteinuria and hypertension in unattended cases. Blood biomarkers for pre-eclampsia are lacking. In this study, microarray gene expression data from peripheral blood of pre-eclampsia women was analyzed. In our study we developed a combined network approach for hub node prediction regulated by transcription factors and microRNAs corresponding to pre-eclampsia. Differentially expressed genes (DEGs) interaction map was constructed using STRING database. JUN, RPL35, NDUFB2, ATP5I, UQCRQ, COX7C, and FN1 were predicted as potential novel hub genes. Pathway analysis showed metabolic pathways, cytokine signaling in the immune system, Wnt, and MAPK signaling pathways involvement in pre-eclampsia. Regulatory network analysis showed that transcription factors JUN and STAT1 were connected with hub nodes, and microRNAs (miRNAs) like hsa-miR-26b-5p and hsa-miR-155-5p. In conclusion, the expression pattern of hub genes, analyzed deciphers a molecular signature for understanding the pathophysiology of pre-eclampsia and prediction of biomarkers for diagnosis.

## 1. Introduction

Pre-eclampsia causes complications in pregnancy and is characterized by uremia, proteinuria, and hypertension (Villar et al., 2003). Almost 8% of all pregnancies are reported to be affected by this gestation-associated syndrome (Li, and Fang, 2019). The early detection of pre-eclampsia is challenging because it shows variable clinical presentations. Two clinical presentations are reported for pre-eclampsia pathophysiology: the first is linked to a decrease in perfusion of the placenta, platelet aggregation, and dysfunction of the endothelium, whereas the second is associate with the immunological abnormalities (Eiland et al., 2012; Redman, and Sargent, 2010). Pre-eclampsia may result in haemolysis, eclampsia, thrombocytopenia, increased serum level of liver enzymes, premature birth, restriction of intrauterine growth, and perinatal fetal death (Li, and Fang, 2019). However, the molecular mechanism and signaling pathways involvement in the pathophysiology of pre-eclampsia is not fully understood. Early detection of pre-eclampsia is essential for effective clinical management. Recent studies have been shown that angiogenic factors like soluble endoglin, placental growth factor (PLGF), soluble vascular endothelial growth factor receptor (sVEGF-R, known as sFLT1) and circulating CD41+/AnnexinV+ microparticles are potential predictive biomarkers of pre-eclampsia. However, these biomarkers were not consistent in predicting the pre-eclampsia (Kleinrouweler et al., 2012; Levine et al., 2006; McElrath et al., 2012).

Recent advancements in sequencing technologies and bioinformatics approaches (Li, and Fang, 2019; Liu et al., 2019). have provided newer and robust tools for the biomarker identification of various disorders (Drucker, and Krapfenbauer, 2013; Parra Cordero et al., 2013). Network modeling techniques are capable of improving and integrating the possibility of disease state complexity and causal factors (Guttula et al., 2018). Network analysis approaches have also provided unique and practical strategies for the prediction of the diseases at an early stage and drug designing (Suresh, and Ashok, 2018). In recent years, several studies have applied microarray technology for studying placental biopsies to identify the pre-eclampsia biomarkers (Founds et al., 2012). A few studies have also been published on the analysis of blood transcriptome in pre-eclampsia patients. (Textoris et al., 2013). Unfortunately, these studies were unable to explain the significance of the blood biomarkers in pre-eclampsia. We hypothesized that the protein-protein interactome (PPI) study might help in understanding the differentially expressed genes (DEGs) and proteins in pre-eclampsia cases, as has been proven to be an advanced and unconventional approach for the prediction of the biomarkers in other diseases (Sriroopreddy, and Sudandiradoss, 2018). Conformational application of statistical methods may also help in the prediction of the hub nodes, regulated by transcription factors and miRNAs corresponding to pre-eclampsia on the retrieved data (Sriroopreddy et al., 2019). It may further assist in retrieving the information to predict the regulatory interactions and to accomplish the goals of therapeutics (SZALLASI, 1999).

Thus, in the current study, an integrative bioinformatics approach was applied to microarray-based gene expression data, from peripheral blood of pre-eclampsia patients, to identify hub genes, and decipher the molecular mechanism of pre-eclampsia pathophysiology by network analysis. The backbone network of DEGs was analyzed using network analysis, and corresponding transcription factors with correlated miRNAs were identified. Besides, potential blood-based molecular markers for the detection of pre-eclampsia were also predicted.

## 2. Materials and Methods

### 2.1. Datasets and their normalization

To identify the molecular markers of pre-eclampsia and the molecular pathways that might be affected during the pre-eclampsia, gene expression datasets from 36 peripheral blood samples of healthy (n= 19) and pre-eclampsia (n=19) patients were retrieved (GEO ID: GSE48424) and analyzed by a variety of bioinformatics tools. The gene expression data were obtained from microarray chips which includes 45,000 probes (Agilent-014850 Whole Human Genome Microarray 4×44K G4112F) and the analysis of expression of One Color Microarray Based Gene chips as described elsewhere (Textoris, Ivorra, Amara, Sabatier, Ménard, Heckenroth, Bretelle, and Mege, 2013). All samples from healthy were considered as controls, whereas pre-eclampsia samples were found as experimental or test groups. The .txt files in raw format were retrieved, and the data were subjected to further analysis by using the RMA algorithm for making them comparable.

### 2.2. Gene expression analysis and clustering

The analyses of gene expression levels and gene ontology (GO) annotation were carried out with GO-Elite software using default options (Zambon et al., 2012). Microarray expression values were reported as log2 values. The log-fold change value was calculated by the geometric subtraction of the pre-eclampsia group from the healthy group for each pair-wise comparison. The max log-fold was calculated as the log2 fold change value among the lowest and highest group expression mean for all conditions within the dataset. The DEGs were identified using a combination of a >0.5-fold change in expression (Textoris, Ivorra, Amara, Sabatier, Ménard, Heckenroth, Bretelle, and Mege, 2013). with a statistical significance of p<0.05 (Moderated t-test) using the Benjamini-Hochberg correction method (Emig et al., 2010). The DEGs were subjected to hierarchical clustering for the identification of the different clusters within the control and test groups (Fendri et al., 2013). The comparison and Gene Set Enrichment Analysis (GSEA) were performed with inbuilt GO-Elite application in AltAnalyze (Emig, Salomonis, Baumbach, Lengauer, Conklin, and Albrecht, 2010)., with FDR adjusted enrichment *p*<0.05, which were considered for further studies.

### 2.3. Lineage analysis and identification of pre-eclampsia specific markers

Cell type/ tissue prediction approaches were applied to identify the variations in the tissue and cell marker genes (Kamath-Rayne et al., 2015). LineageProfiler gene marker database was used, which was derived from the hundreds of distinct tissue and cell markers in the GO-Elite software, and the Fischer-Exact enrichment test with *p*<0.05 was used for analyses. To identify the cell/tissue markers in both pre-eclampsia and healthy samples, we first adopted the gene filtration with an expression level cut off of >90 percentile in all samples, and then highly expressed genes were compared in both pre-eclampsia, and healthy samples for the identification of both common and uniquely expressed genes. Further, the MarkerFinder algorithm in AltAnalyze was used within the datasets for the derivation of putative cell-population-specific markers (Hulin et al., 2019).

### 2.4. Pathway enrichment analysis and construction of PPI network

The DEGs were submitted to DAVID server (Jiao et al., 2012). (https://david.ncifcrf.gov/tools.jsp) for obtaining the gene ontology information. Pathway enrichment analysis was executed using the KOBAS tool (Xie et al., 2011). (http://kobas.cbi.pku.edu.cn/) with p< 0.05 to test the statistical significance. The gene list was then introduced to the STRING database (Szklarczyk et al., 2016). (https://string-db.org/cgi/input.pl?sessionId=KRkiiBZSNz5Y) (version: 11.0), to attain the PPI map. The Cytoscape (Shannon et al., 2003). (version: 3.7.2) software was used to analyze and visualize the network topology indexes.

### 2.5. Analysis of protein-protein interaction (PPI) network and prediction of hub nodes/genes

Interaction networks of DEGs, generated from the STRING database, were analyzed using *NetworkAnalyzer* (Saito et al., 2012). in Cytoscape. The topological network parameters were taken into account to accomplish the study of the PPI network. These parameters included betweenness centrality, closeness centrality, degree distribution, and the shortest path distribution. Degree of the node is defined as number of connections linked to it and it is represented as how one node is connected to each other.

Hub genes within the network were identified using *cytoHubba* (Chin et al., 2014). in the Cytoscape. Hub genes were identified by node ranking in the networks, based on the topological analysis methods such as degree, Maximum Neighborhood Component, and Maximal Clique Centrality.

### 2.6. Validation of predicted hub nodes/genes by coexpression analysis

The FpClass (http://dcv.uhnres.utoronto.ca/FPCLASS/) bioinformatics tool was used to validate the hub genes identified in the above analysis. This tool helps to find the interaction and proteins that are co-expressed with the genes submitted. The analysis was performed by considering two individual scores i.e. Gene coexpression score and Network topology score (Kotlyar et al., 2015).

### 2.7. Prediction of transcription factors and miRNAs targeting the hub nodes

The transcription factors targeting the hub nodes were predicted using the i-Regulon (Verfaillie et al., 2014). in Cytoscape. The corresponding transcription factors and hub node pairs with Normalized Enrichment Score (NES)>4 were selected for the analysis. In the next step, miRNAs targeting the transcription factors were predicted using the miRTarbase database (Hsu et al., 2011).. miRNAs, which are common for the transcription factors, were filtered out.

## 3. Results

### 3.1. Normalization of gene expression datasets

Multiple primary quality control (QC) plots were generated for evaluating the quality of the samples and overall technical correlation compared with other samples within the dataset. The distribution of probe intensities (Figure 1) varied from 0 to 2500 observations between pre-eclampsia and healthy samples.

**Figure 1:**
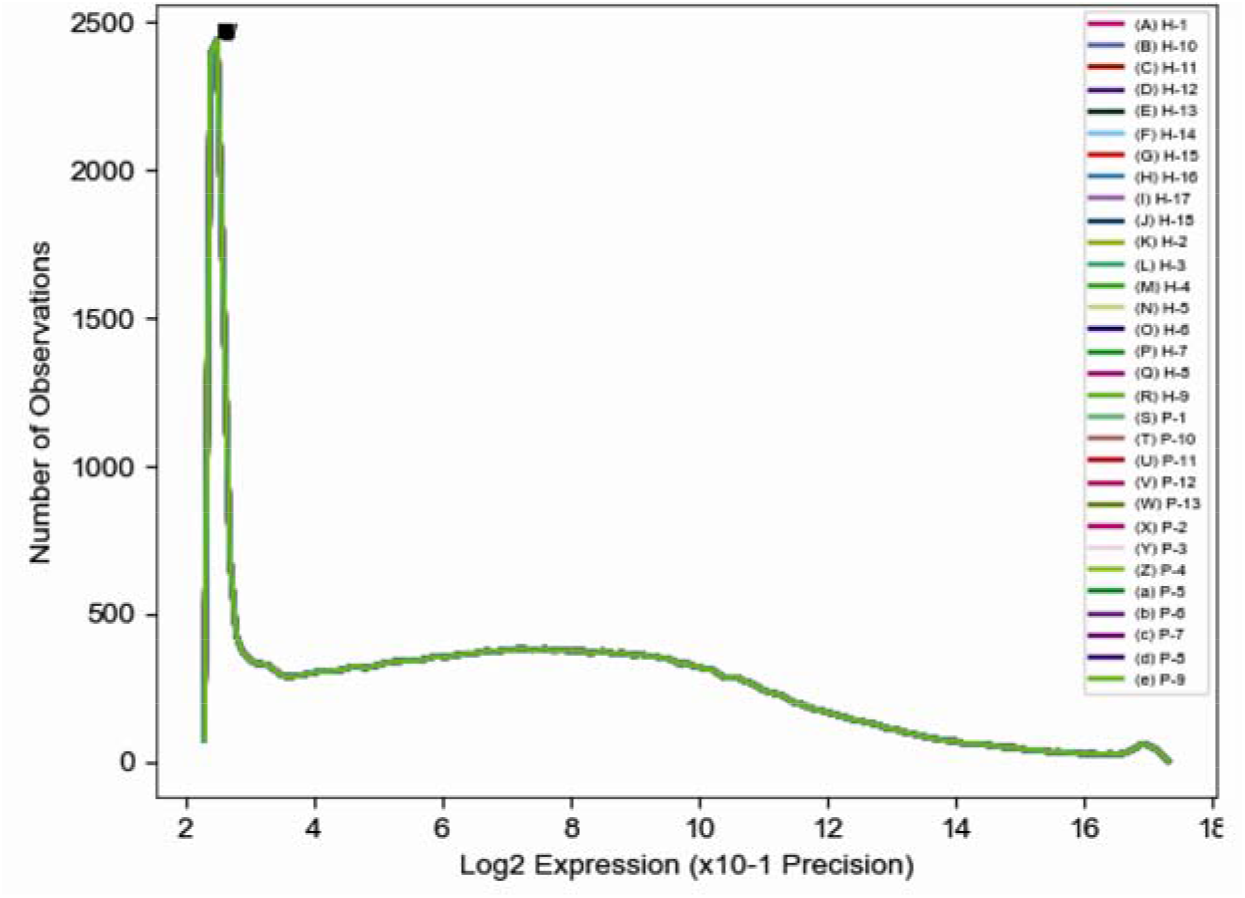
Distribution of microarray probe intensities in peripheral blood samples of pre-eclampsia (n=19) and healthy (n=19) women. H: Healthy women; P: Pre-eclampsia women. Numbers against H and P indicate the sample number.

### 3.2. Differential expression, clustering, and principal component analysis

In comparison to healthy samples, 965 genes were found to be expressed differentially in pre-eclampsia samples. Among these, 400 were found to be up-regulated (Table 1), and 565 found to be down-regulated in pre-eclampsia samples (Table 1). To study the relationship between the gene expression pattern in various samples, the clustering of DEGs was used (Figure 2). Both healthy and pre-eclampsia groups were found to cluster separately upon hierarchical clustering. Principal component analysis (PCA) of the two groups revealed that all pre-eclampsia samples were sharing one component, while all healthy samples were sharing other principal components (Figure 3).

**Table 1:** Top 10 up- and down-regulated genes in the peripheral blood of pre-eclampsia patients

| Gene Symbol | Fold Change | P value |
| --- | --- | --- |
| **Up-regulated Genes** |  |  |
| OLAH | 2.09 | 3.04E-3 |
| VSIG4 | 2.01 | 5.75E-5 |
| FN1 | 1.73 | 1.01E-2 |
| MACROD2 | 1.61 | 1.06E-4 |
| HRK | 1.61 | 1.02E-2 |
| A_23_P8812 | 1.52 | 1.58E-3 |
| CA12 | 1.45 | 1.55E-3 |
| A_32_P156786 | 1.43 | 1.24E-2 |
| LOC283454 | 1.41 | 1.89E-2 |
| DNAH7 | 1.40 | 1.64E-2 |
| **Down-regulated Genes** |  |  |
| B3GNT2 | -0.58542 | 1.26E-2 |
| ITGB2 | -0.58547 | 2.25E-2 |
| PDE8B | -0.58591 | 1.48E-2 |
| EFHB | -0.58619 | 2.07E-3 |
| DDX3X | -0.58666 | 5.08E-3 |
| THC1954314 | -0.58687 | 2.64E-4 |
| APC | -0.58700 | 4.04E-2 |
| CNKSR3 | -0.58769 | 7.15E-3 |
| MFSD14B | -0.58778 | 8.65E-3 |
| A_24_P247117 | -0.58796 | 2.37E-2 |

**Figure 2:**
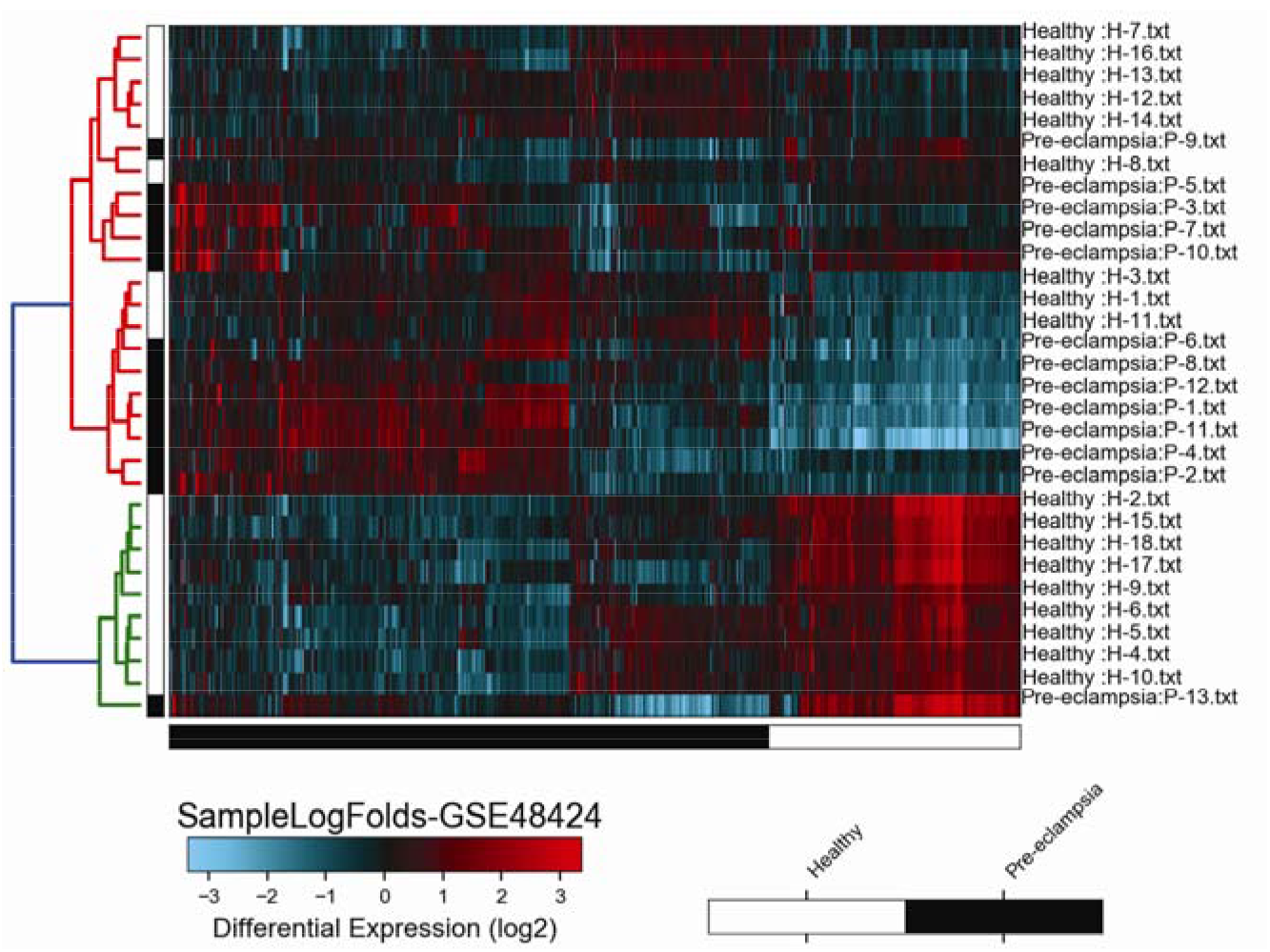
Heatmap showing the log-fold changes and hierarchal clustering of various genes expressed in the peripheral blood of pre-eclampsia and healthy women. H: Healthy women; P: Pre-eclampsia women. Numbers against H and P indicate the sample number.

**Figure 3:**
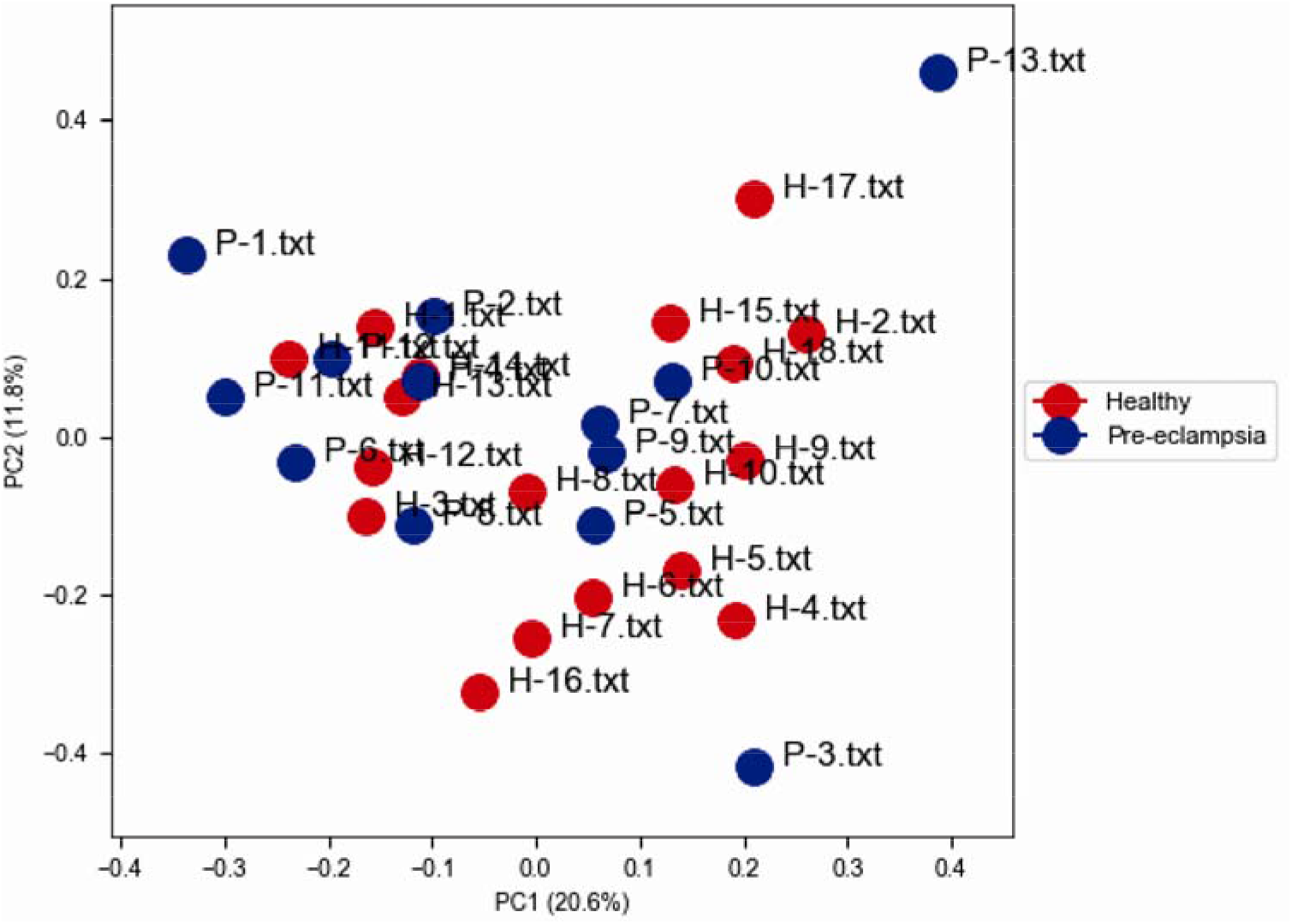
Principal component analysis (PCA) of pre-eclampsia and healthy samples sharing different principal components. Red spheres represent pre-eclampsia samples whereas blue spheres represent healthy samples.

### 3.3. Lineage analysis

The result of lineage analysis is shown as a lineage correlation heat map (Figure 4). The lineage correlation heat map revealed that both healthy and pre-eclampsia Z score sharing was high with neutrophils, blood, spleen, and dendritic cells indicating that these genes are highly expressed in these tissues.

**Figure 4:**
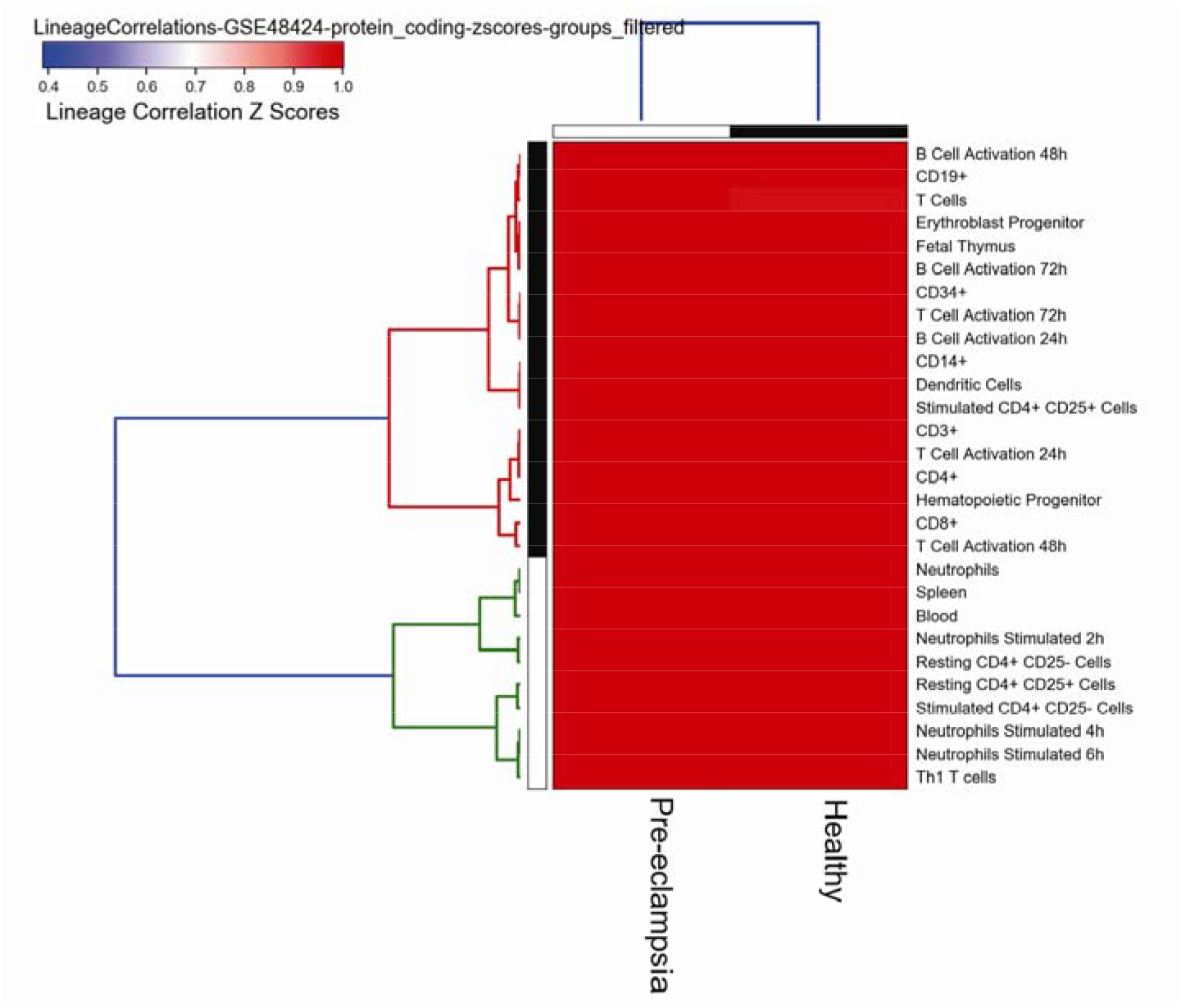
Lineage correlation analysis of genes, expressed in pre-eclampsia and healthy samples, with other cell and tissue types.

### 3.4. Marker gene identification

A total of 60 pre-eclampsia specific genes were identified by Pearson correlation coefficient analysis (Figure 5). GPER1, RPS15AP12, C1QA, RPS26P39, and RPL26L1 are predicted as markers for pre-eclampsia and were not present in the top 10. The top 10 genes (Table 2), identified as pre-eclampsia markers, included MTMR9, CCDC127, ADAM17, and PDDC1. On the other end, the top 10 genes (Table 2), identified as marker genes in healthy samples, included RNF38, PAPD4, SGPP1, and HIAT1. Table 2 includes only the top 10 marker genes for both pre-eclampsia and healthy samples.

**Figure 5:**
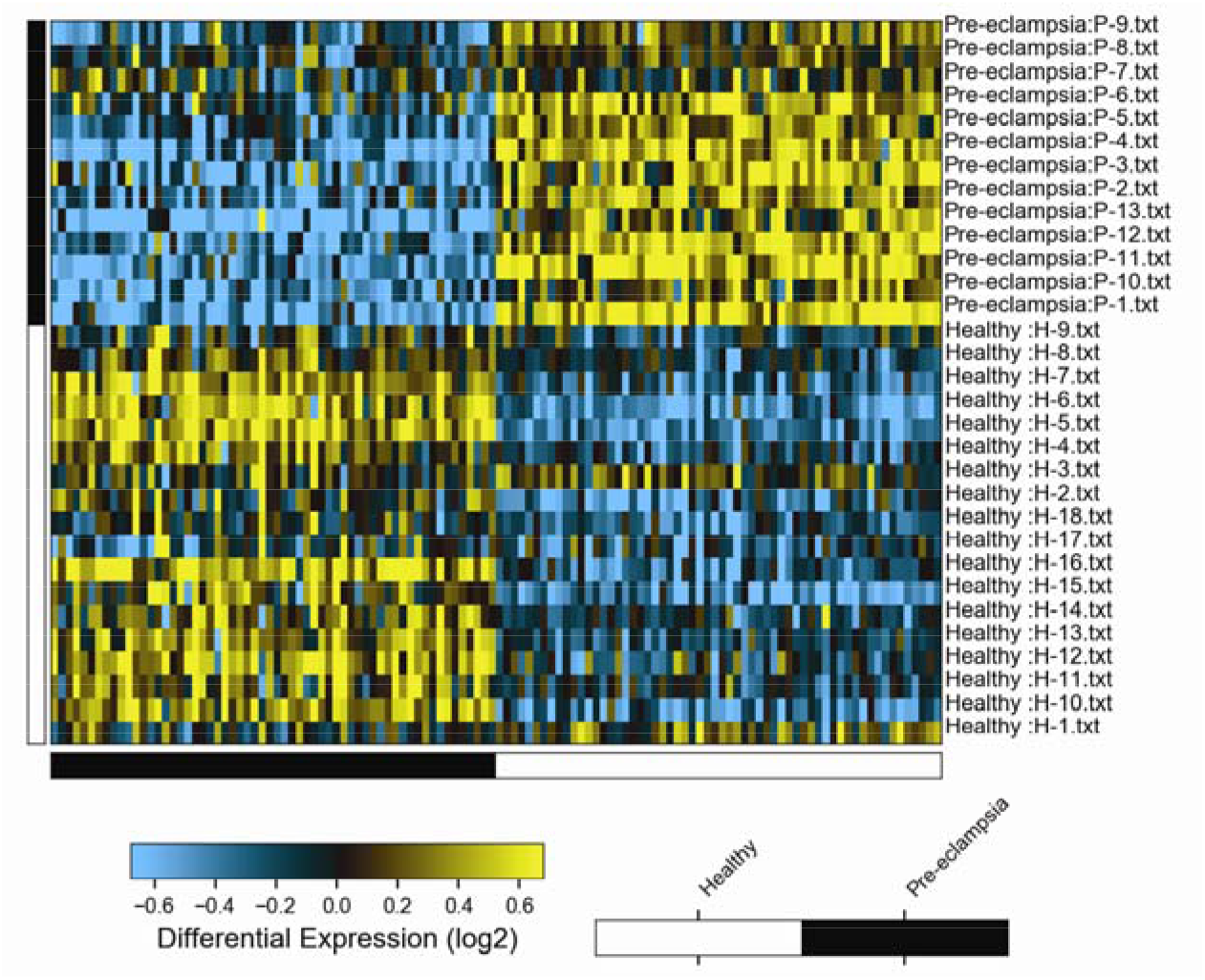
ICGS analysis of transcriptomes for finding markers of pre-eclampsia and healthy samples. H: Healthy women; P: Pre-eclampsia women. Numbers against H and P indicate the sample number.

**Table 2:**
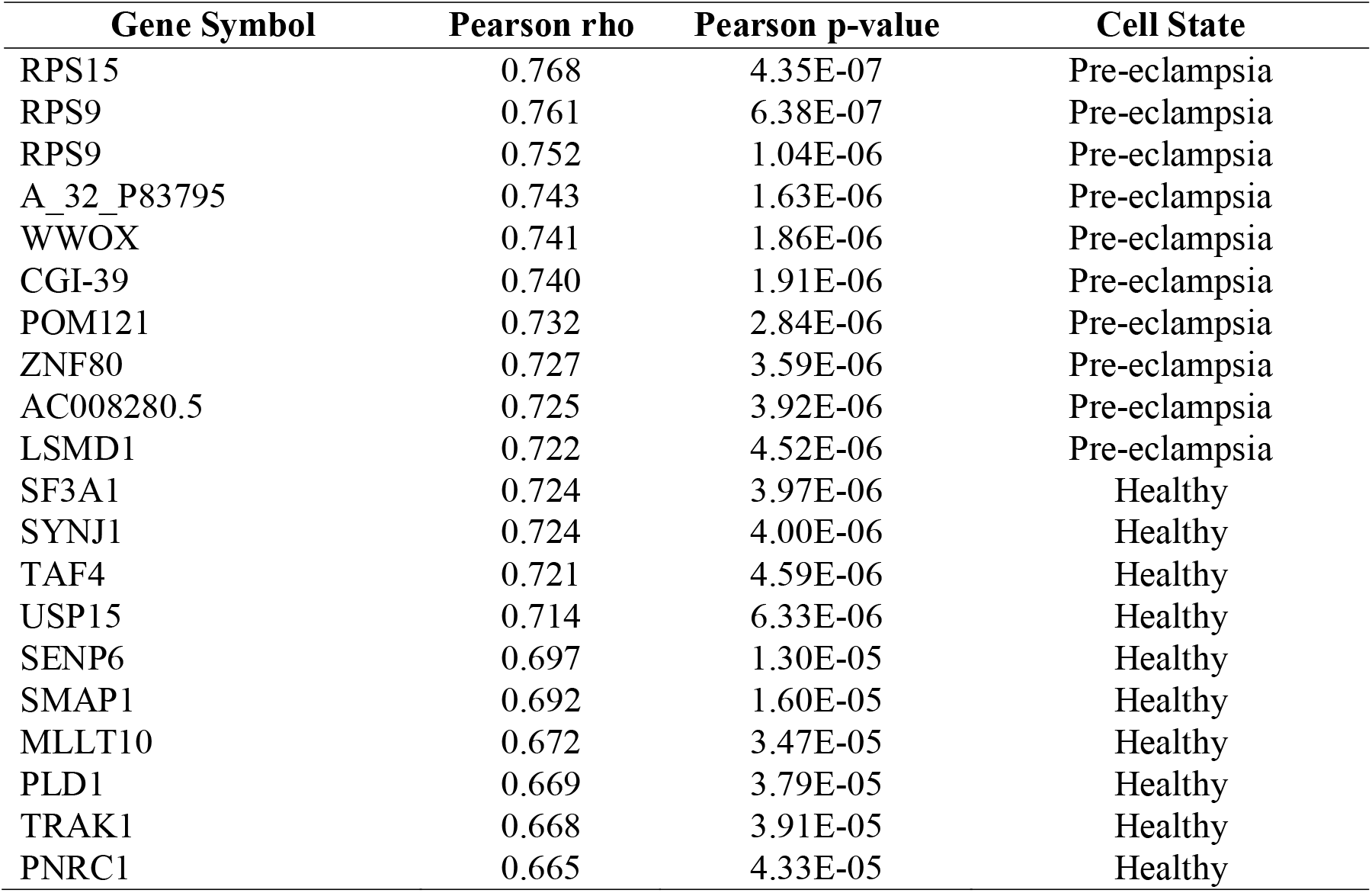
Potential marker genes for detection of pre-eclampsia in peripheral blood samples

### 3.5. Gene ontology analysis

The functional annotation of DEGs was executed in the DAVID functional annotation server and it predicted the specific biological functions like cell communication, cell cycle, and cell differentiation. Various genes were also associated with essential cellular functions like immune regulation, cell death, and hemopoiesis, etc.

### 3.6. Pathway enrichment analysis

The pathway enrichment analysis was performed using the KOBAS tool. The identified DEGs such as JUN, RPS21, FN1, ADAM17, and NOS2 were enriched significantly in the immune system, metabolic pathways, cytokine signaling in the immune system, Wnt signaling pathway, Jak-STAT, MAP kinase, Jak-STAT, Wnt, Notch and NF-kappa signaling pathways (Table 3).

**Table 3:**
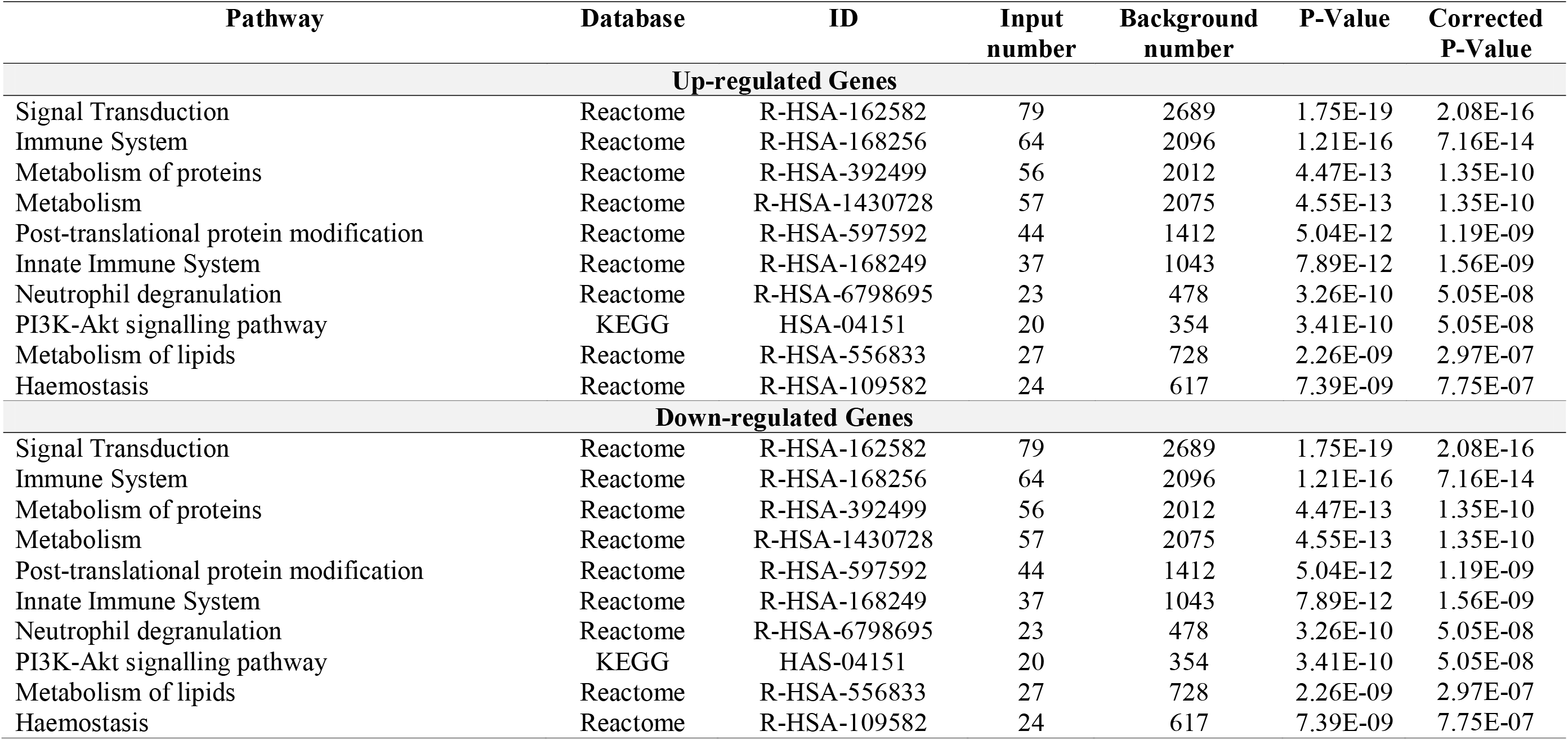
Pathway enrichment analysis of up- and down-regulated genes in the peripheral blood of pre-eclampsia patients

### 3.7. PPI network and prediction of hub nodes

The PPI interactome was generated using the STRING database for both up- and down-regulated genes and was visualized in Cytoscape. The identified backbone networks comprised 400 up-regulated (Figure 6A) and 565 down-regulated (Figure 6B) genes with an average local clustering coefficient of 0.377 and 0.373, respectively. The networks were further analyzed for their topological parameters by *NetworkAnalyser*. The pictorial representation of the topological parametres for both up-regulated and down-regulated networks are shown in Figures 7 and 8, respectively. On the basis of topological parameters, such as degree, path length, closeness, and betweenness (Table 4), the hub genes were selected. High confidence was seen in the network’s clustering coefficient. Top 20 nodes (Table 5) are highly connected with degree >13. The up-regulated hub genes (Figure 9A) include JUN, RPL35, CD19, ATP5I, SLIRP, etc. whereas the down-regulated hub genes (Figure 9B) included VEGFA, CAV1, CCR1, PLD1, etc.

**Table 4:** Topological parameters of the up- and down-regulated PPI network.

| **Topological parameters** | **Comprehended values** |
| --- | --- |
| **Up-regulated genes** |  |
| Number of nodes | 188 |
| Number of edges | 458 |
| Clustering co-efficient | 0.290 |
| Network density | 0.026 |
| Characteristic path length | 4.398 |
| Average number of neighbours | 4.872 |
| **Down-regulated genes** |  |
| Number of nodes | 388 |
| Number of edges | 886 |
| Clustering co-efficient | 0.186 |
| Network density | 0.012 |
| Characteristic path length | 4.502 |
| Average number of neighbours | 4.567 |

**Table 5:** Centrality values of top 10 up-regulated and down-regulated genes in pre-eclampsia.

| Node Name | Degree | Closeness | Betweenness |
| --- | --- | --- | --- |
| **Up-regulated Genes** |  |  |  |
| FN1 | 23 | 76.48 | 8122.09 |
| COX7C | 22 | 59.96 | 1125 |
| RPL9 | 21 | 60.91 | 748.3 |
| RPL27 | 20 | 60.57 | 528.93 |
| CD19 | 18 | 67.17 | 3747.98 |
| RPS21 | 18 | 58.77 | 263.35 |
| UQCRQ | 17 | 57.16 | 950 |
| PFDN5 | 16 | 65.25 | 7594.84 |
| RPL26L1 | 16 | 55.42 | 191.29 |
| RPL35A | 16 | 57.99 | 169.22 |
| **Down-regulated Genes** |  |  |  |
| EGF | 40 | 158.48 | 22468.02 |
| VEGFA | 37 | 152.76 | 17496.05 |
| FPR2 | 30 | 138.21 | 4930.84 |
| IL1B | 29 | 145.54 | 9703.03 |
| TLR4 | 27 | 143.17 | 7822.55 |
| ITGB2 | 25 | 139.44 | 8186.5 |
| CAV1 | 22 | 142.32 | 11786.09 |
| RAB11A | 21 | 134.49 | 7733.81 |
| SNAP23 | 21 | 134.87 | 5636.33 |
| MGAM | 20 | 128.41 | 11214.23 |

**Figure 6:**
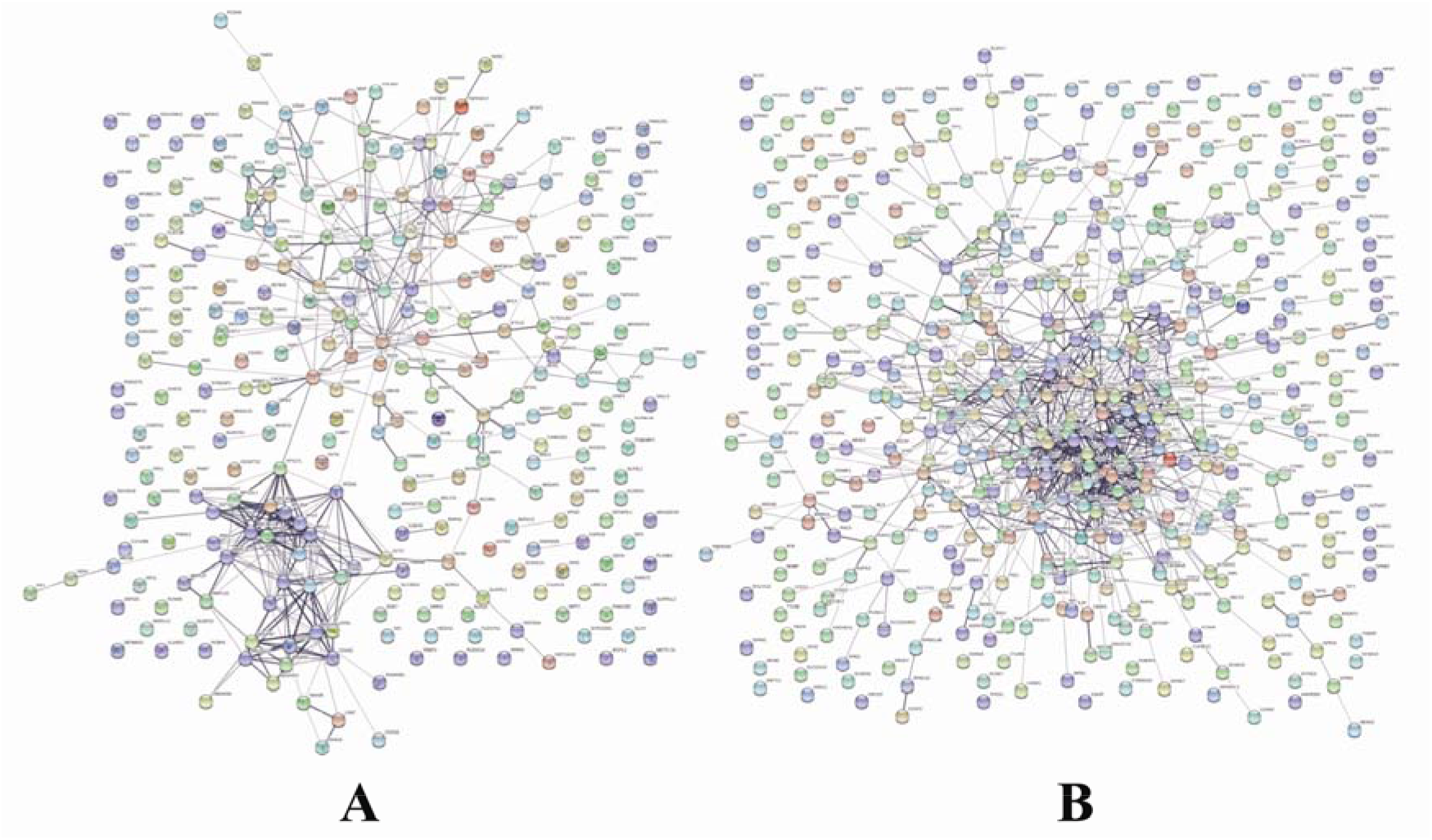
Backbone network of up-regulated (A) and down-regulated (B) genes among healthy and pre-eclampsia samples. The spheres represent the nodes whereas the lines indicate the edges.

**Figure 7:**
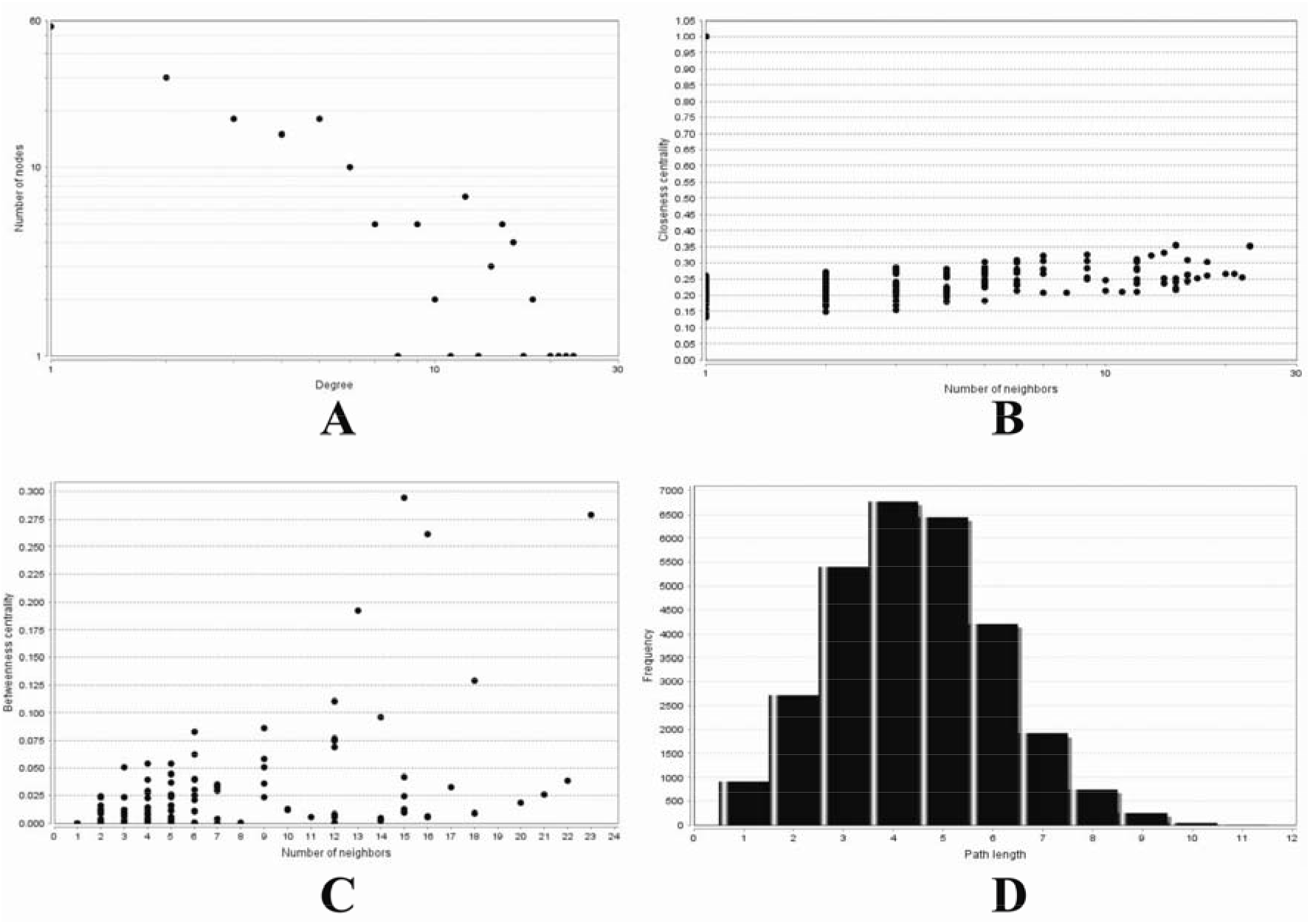
Topological parameters of the backbone network for up-regulated differentially expressed genes. A: Degree distribution; B: Closeness centrality; C: Betweenness centrality; D: Shortest path distribution.

**Figure 8:**
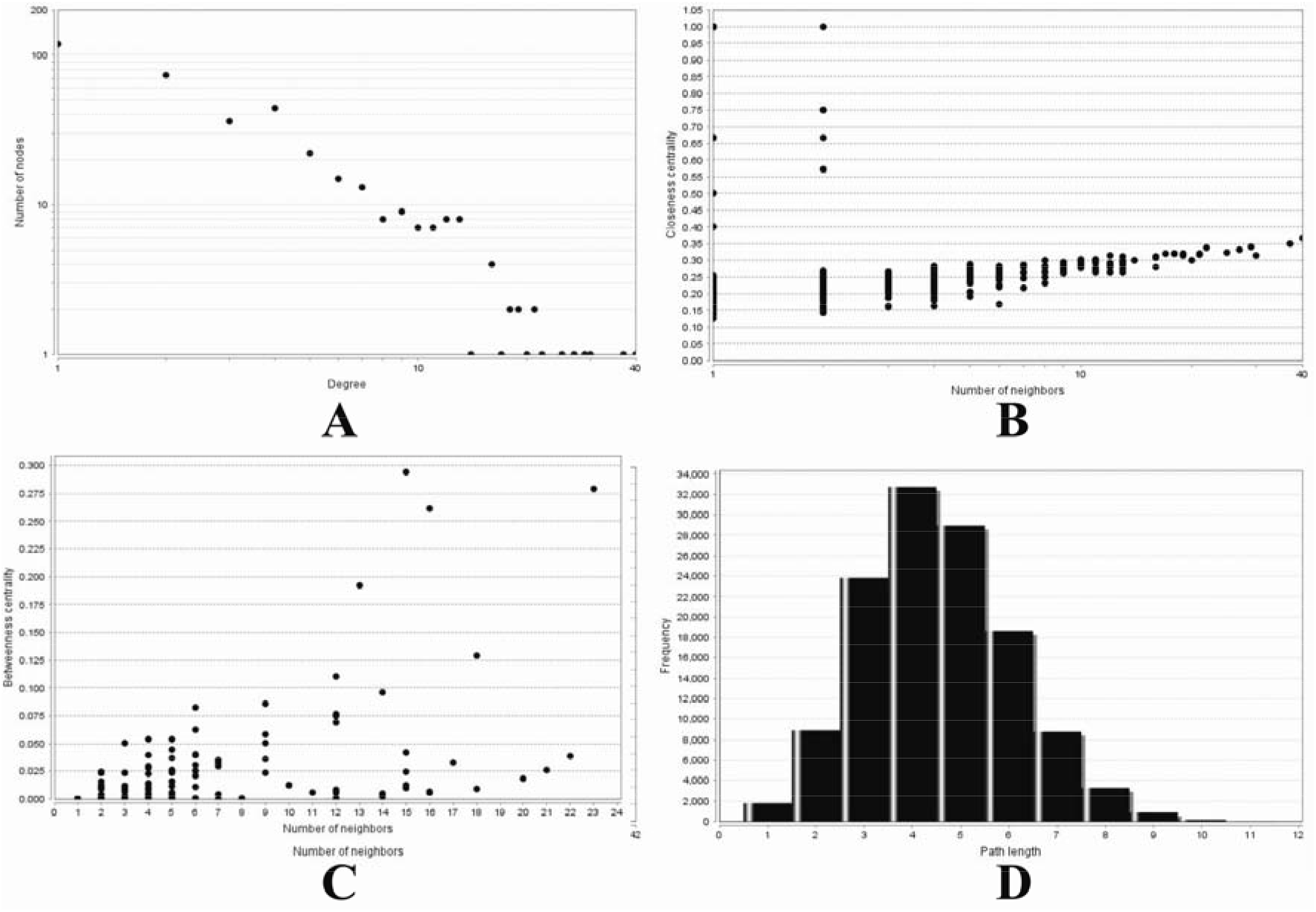
Topological parameters of the backbone network for down-regulated differentially expressed genes. A: Degree distribution; B: Closeness centrality; C: Betweenness centrality; D: Shortest path distribution.

**Figure 9:**
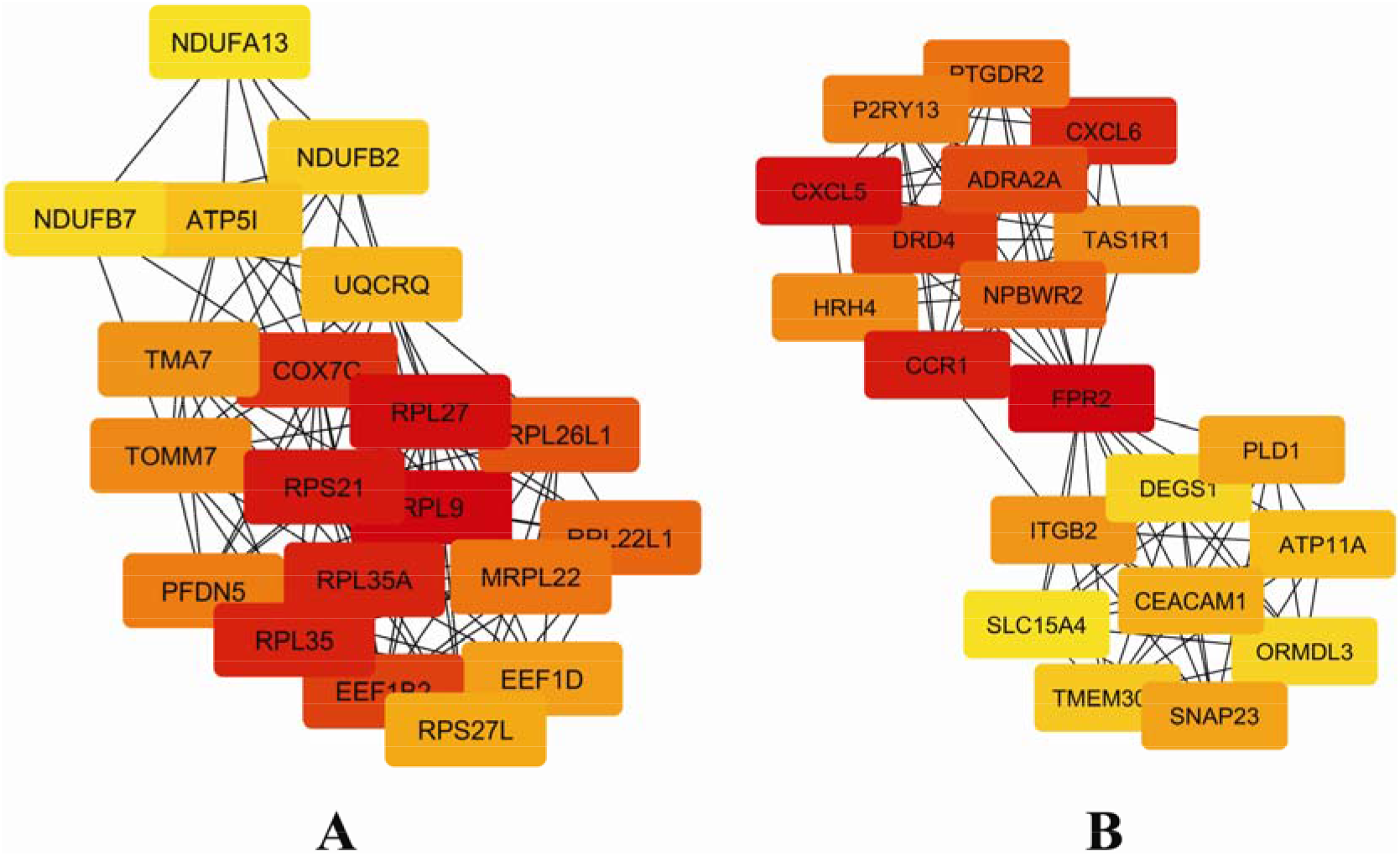
Hub node network for top 20 (A) up-regulated genes and (B) down-regulated genes.

### 3.8. Validation of hub genes by co-expression analysis

The FpClass tool was used to validate the identified hub genes by studying their coexpression and their interacting genes. The partner genes were picked on the basis of highest total score. Among these genes, the PLCG2 gene showed the top-most network topology score of about 0.6887 and showed interaction with the CD19 gene to be stable.

### 3.9. Transcription factors and related recognition of miRNA

Transcription factors that are linked with hub genes were identified using i-Regulon. This analysis resulted in the prediction of potential transcription factors that included ESR2, JUN, ATF1, STAT1, HNF1A, STAT6 with NES score of higher than 4. A total of 24 transcription factors were identified. A Total of six miRNAs (Table 6), which are common, were linked to at least two transcription factors. miRNAs such as hsa-miR-155-5p and hsa-miR-203a-3p were found to target both JUN and STAT1. It was found that almost all six miRNAs targeted the JUN transcription factor. A regulatory network was constructed with the information of miRNAs, genes, and transcription factors (Figure 10) wherein hsa-miR-26b-5p and hsa-miR-155-5p were noticed to be most outstanding ones.

**Table 6:** List of miRNAs associated with the common transcription factors from miRTarbase.

| Term | Overlap | P-value | Adjusted P-value | Odds Ratio | Combined Score |  |
| --- | --- | --- | --- | --- | --- | --- |
| hsa-miR-155-5p | 6/903 | 5.57E-04 | 0.164 | 5.537 | 41.491 | JUN;CBFB;STAT1; |
| hsa-miR-203a-3p | 3/307 | 5.71E-3 | 0.451 | 8.143 | 42.066 | JUN;STAT1;NR2C |
| hsa-miR-140-5p | 2/100 | 6.36E-3 | 0.479 | 16.666 | 84.299 | STAT1;ESR2 |
| hsa-miR-93-5p | 5/1220 | 1.346E-2 | 0.641 | 3.415 | 14.712 | JUN;POLR2A;E2F |
| hsa-miR-26b-5p | 6/1872 | 2.055E-2 | 0.765 | 2.670 | 10.375 | JUN;FOXD2;MEF2 |
| hsa-miR-150-5p | 3/534 | 2.526E-2 | 0.826 | 4.681 | 17.221 | STAT1;TP53;ESR2 |

**Figure 10:**
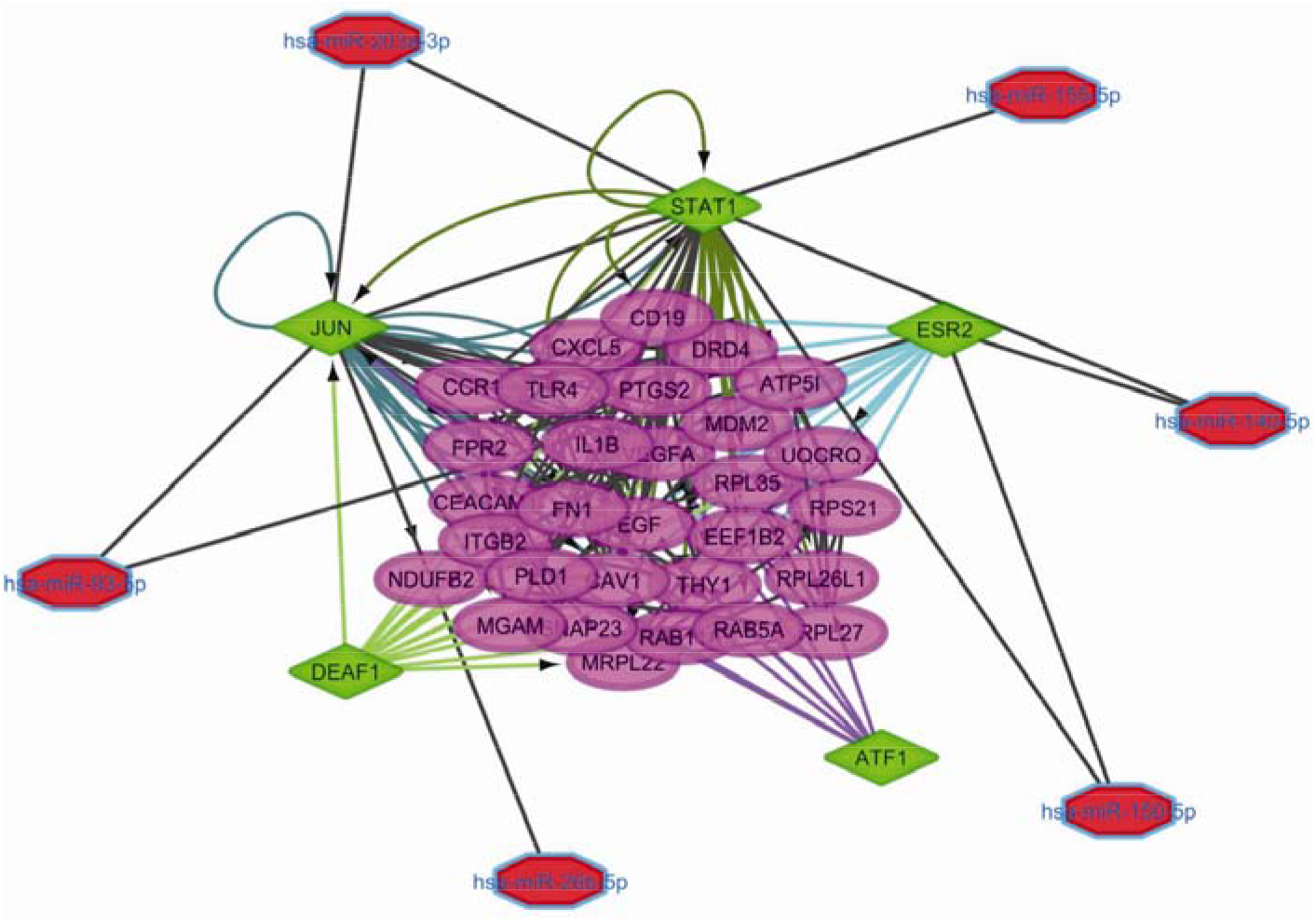
The transcription regulatory network depicting the hub genes connected to the associated transcriptional factors and the correlated miRNAs to the transcription factors in pre-eclampsia. The central bigger nodes are the hub genes, the transcriptional factors are colored in blue and the miRNAs in the outer circle are numbered and colored in gr

## 4. Discussion

In the current study, an computational analysis of publicly available microarray data was performed for the prediction of hub genes/nodes integrating with transcription factors and miRNAs as potential blood biomarkers for pre-eclampsia. The gene ontology and pathway enrichment of DEGs suggested their enrichment in disease-related biological processes and pathways. From the observed results, the up-regulated genes were also significantly enriched in the metabolic pathways. According to our results, various signalling pathways such as the MAP kinase, Jak-STAT, NOTCH and NF-kappa were found to be affected in the pre-eclampsia. The components of Notch signaling were earlier reported to be affected by the disorders of pregnancy, like pre-eclampsia, which plays a crucial part in the placental developmental processes (Haider et al., 2017).

From the analysis, 39 hub genes/nodes were predicted as potential blood biomarkers for pre-eclampsia. Genes such as RPL35, RPL35A, and RPL27 observed in our study were reported to be involved in the biogenesis of ribosome, suggesting that the characterization of pre-eclampsia is due to change in the machinery of the translation (Textoris, Ivorra, Amara, Sabatier, Ménard, Heckenroth, Bretelle, and Mege, 2013). PTGS2, one of the hub nodes in our study, was downregulated in pre-eclampsia. Published studies have reported that PTGS2 gets associated with the miR-144-3p and participates in the pre-eclampsia pathogenesis (Hu et al., 2019). Epidermal growth factor (EGF) predicted as a hub node in our study was downregulated in pre-eclampsia (Kupsamy et al., 2019). and may be a promising biomarker to diagnose pre-eclampsia. It has been suggested that EGF is dysregulated in the placenta, and disruption of the EGF signaling system contributes to aberrant trophoblast development in pre-eclampsia (Armant et al., 2015). VEGFA is a hub node predicted in our study and is found to get downregulated in pre-eclampsia. It was earlier reported to be a direct target for miR-199a-5p, and its expression is decreased in pre-eclampsia (Mei et al., 2019).

Marker genes that were identified using the MarkerFinder algorithm were compared with the hub nodes to evaluate their expression in pre-eclampsia and healthy samples. According to the results, three hub genes, such as PI3, CCR1, and RAB5A, were predicted as markers in the healthy samples, and these genes are downregulated in pre-eclampsia in our study. Novel hub genes were identified from the analysis according to pathway enrichment studies, which showed that FN1 participates in the immune system and cytokine signaling pathways. On the other hand, novel up-regulated genes such as NDUFB2, ATP5I, UQCRQ, and COX7C were found to participate in metabolic pathways and may get associated with the serum levels in pre-eclampsia. Another new gene, JUN, found to participate in the NOTCH signaling, and its dysregulation may lead to pre-eclampsia. Coexpression partner validation results of our study suggest that PLCG2 has the highest network topology score that interacts with CD19, and it is reported as an indicator for pre-eclampsia in previous studies (Jensen et al., 2012).

Transcription factors that are primary regulators of gene expression in pre-eclampsia are ESR2, ESR1, RARB, CEBPD, CEBPB, CEBPG, etc. Transcription factors of hub genes identified in our study stimulate the genes by post translational mechanisms like sumoylation, dephosphorylation, and phosphorylation (Sriroopreddy, Sajeed, Raghuraman, and Sudandiradoss, 2019). Transcription factors, such as JUN, STAT1, ESR2, and ATF1, were linked with the various hub genes and controlled their gene expression. Studies have reported that JUN activates the signaling pathway related to C-Jun-dependent kinases, and it was down-regulated in pre-eclampsia (Hannke-Lohmann et al., 2000). Phosphorylation of STAT1 occurs by dimer formation and activates the Janus tyrosine kinases (JAKs) during the initiation by IFN-γ (Aaronson, and Horvath, 2002). Activation of endothelial cells and excess inflammation are considered as a features of pre-eclampsia by the induction of IFN-γ/STAT1 signaling pathway (Liu et al., 2016).

The above-discussed hub genes and transcription factors we observed by pathway analysis were enriched significantly in immune system-related pathways, for example, JUN and STAT1. By compiling both experimental and clinical evidence, it was clear that there is an involvement of the immune system in the pathogenesis of pre-eclampsia. The changes in the immune system and T-Cell alterations were linked with the origin of pre-eclampsia. This might suggest that a lack of balance in the immune system may worsen the state of the inflammation in pre-eclampsia.

In summary, the results obtained by our analysis establish the differential expression of corresponding hub nodes and their mechanism. Discussed hub nodes could act as potential biomarkers for pre-eclampsia. The role of every hub node and its partners can outweigh the understanding of transition of pre-eclampsia and commits towards improved support to reach the specific ongoing needs in order to develop the efficiency for leading treatment selection and differential diagnosis. In conclusion, the expression pattern of hub genes, analyzed by our approach, may be considered as a molecular signature to decipher the pathophysiology of pre-eclampsia and genes such as JUN, RPL35, RPL35A and PTGS2 etc along with pathways like immune system related, NOTCH signaling etc could also offer as potential biomarkers for pre-eclampsia prognosis and diagnosis.

## Acknowledgements

The authors acknowledge Indian council of Medical Research for providing the funding, Department of Biotechnology for providing the Infrastructure facility.

## Notes

**Funding Details:** The work was supported by the Indian Council of Medical Research under Grant number ISRM/11(29)/2017 and Department of Biotechnology under grant number BT/BI/25/051/2015 dt: 21.05.2014

### Competing Interest Statement

The authors have declared no competing interest.

