## Supplementary Data for "Protein-protein interaction map in pre-eclampsia through the integration of hub genes, transcription factors and microRNAs"

**Table S1:** Gene ontology analysis of the DEGs

| **Category** | **Term** | **Count** | **%** | **PValue** | **Genes** |
| --- | --- | --- | --- | --- | --- |
| GOTERM_BP_DIRECT | GO:0006412~translation | 12 | 3.351955 | 0.001611 | RPL35A, MRPL22, RPL41, RPL9, RPL26L1, RPL35, RPL27, RPS27L, RPL22L1, RPS21, ZNF525, MRPL43 |
| GOTERM_BP_DIRECT | GO:0032981~mitochondrial respiratory chain complex I assembly | 6 | 1.675978 | 0.002575 | NDUFB7, NDUFAF8, NDUFA13, NDUFAF2, NDUFB1, NDUFB2 |
| GOTERM_BP_DIRECT | GO:0006614~SRP-dependent cotranslational protein targeting to membrane | 7 | 1.955307 | 0.002973 | RPL35A, RPL41, RPL9, RPL26L1, RPL35, RPL27, RPS21 |
| GOTERM_BP_DIRECT | GO:0008284~positive regulation of cell proliferation | 16 | 4.469274 | 0.004861 | FLT3, MZB1, LIFR, SOX4, GPER1, GDNF, GLI1, AKR1C3, EPCAM, SPDYA, HIPK2, PDGFRA, MDM2, ADAM17, EMP2, FN1 |
| GOTERM_BP_DIRECT | GO:0002181~cytoplasmic translation | 4 | 1.117318 | 0.006084 | RPL35A, RPL9, RPL26L1, RPL22L1 |
| GOTERM_BP_DIRECT | GO:0071560~cellular response to transforming growth factor beta stimulus | 5 | 1.396648 | 0.006268 | CAV1, CLEC3B, XCL1, PPARGC1A, WWOX |
| GOTERM_BP_DIRECT | GO:0019083~viral transcription | 7 | 1.955307 | 0.00699 | RPL35A, RPL41, RPL9, RPL26L1, RPL35, RPL27, RPS21 |
| GOTERM_BP_DIRECT | GO:0000184~nuclear-transcribed mRNA catabolic process, nonsense-mediated decay | 7 | 1.955307 | 0.0093 | RPL35A, RPL41, RPL9, RPL26L1, RPL35, RPL27, RPS21 |
| GOTERM_BP_DIRECT | GO:0010952~positive regulation of peptidase activity | 3 | 0.837989 | 0.015808 | CAV1, NDUFA13, FN1 |
| GOTERM_BP_DIRECT | GO:0006364~rRNA processing | 9 | 2.513966 | 0.015824 | RPL35A, RPL41, RPL9, RPL26L1, RPL35, NPM3, RPL27, RPS27L, RPS21 |
| GOTERM_BP_DIRECT | GO:0043627~response to estrogen | 5 | 1.396648 | 0.016628 | ARPC1B, GSTM3, CAV1, PPARG, RGS9 |
| GOTERM_BP_DIRECT | GO:0006413~translational initiation | 7 | 1.955307 | 0.017655 | RPL35A, RPL41, RPL9, RPL26L1, RPL35, RPL27, RPS21 |
| GOTERM_BP_DIRECT | GO:0043547~positive regulation of GTPase activity | 16 | 4.469274 | 0.024623 | ARHGEF28, ARHGAP19, AKAP13, S100A10, HERC2, ARHGAP24, GDNF, AMPH, JUN, PDGFRA, RGS9, SYNGAP1, AXIN2, XCL1, XCL2, SRGAP1 |
| GOTERM_BP_DIRECT | GO:1903575~cornified envelope assembly | 2 | 0.558659 | 0.029907 | STX2, DMKN |
| GOTERM_BP_DIRECT | GO:1901558~response to metformin | 2 | 0.558659 | 0.029907 | PPARG, PPARGC1A |
| GOTERM_BP_DIRECT | GO:0071375~cellular response to peptide hormone stimulus | 3 | 0.837989 | 0.035943 | CAV1, MDM2, GPER1 |
| GOTERM_BP_DIRECT | GO:0034614~cellular response to reactive oxygen species | 3 | 0.837989 | 0.035943 | AKR1C3, SLC8A1, PDGFRA |
| GOTERM_BP_DIRECT | GO:0006123~mitochondrial electron transport, cytochrome c to oxygen | 3 | 0.837989 | 0.035943 | COX7B, COX7C, COX6C |
| GOTERM_BP_DIRECT | GO:0060070~canonical Wnt signaling pathway | 5 | 1.396648 | 0.036681 | WNT3, NR4A2, SOX4, UBE2B, GLI1 |
| GOTERM_BP_DIRECT | GO:0006120~mitochondrial electron transport, NADH to ubiquinone | 4 | 1.117318 | 0.037473 | NDUFB7, NDUFA13, NDUFB1, NDUFB2 |
| GOTERM_BP_DIRECT | GO:0006974~cellular response to DNA damage stimulus | 8 | 2.234637 | 0.037832 | BATF, SPDYA, TP53TG1, NUAK1, MACROD2, RPS27L, HERC2, UBE2B |
| GOTERM_BP_DIRECT | GO:1904637~cellular response to ionomycin | 2 | 0.558659 | 0.044525 | ZNF683, PPARGC1A |
| GOTERM_BP_DIRECT | GO:0043524~negative regulation of neuron apoptotic process | 6 | 1.675978 | 0.049585 | JUN, HIPK2, NR4A2, SYNGAP1, GDNF, PPARGC1A |
| GOTERM_BP_DIRECT | GO:0030511~positive regulation of transforming growth factor beta receptor signaling pathway | 3 | 0.837989 | 0.050211 | CDKN2B, HIPK2, ADAM17 |
| GOTERM_BP_DIRECT | GO:0051899~membrane depolarization | 3 | 0.837989 | 0.054048 | CAV1, JUN, CACNG2 |
| GOTERM_BP_DIRECT | GO:0031018~endocrine pancreas development | 3 | 0.837989 | 0.057985 | ANXA1, SOX4, RFX3 |
| GOTERM_BP_DIRECT | GO:2001238~positive regulation of extrinsic apoptotic signaling pathway | 3 | 0.837989 | 0.057985 | CAV1, GPER1, WWOX |
| GOTERM_BP_DIRECT | GO:0002328~pro-B cell differentiation | 2 | 0.558659 | 0.058924 | FLT3, SOX4 |
| GOTERM_BP_DIRECT | GO:0051281~positive regulation of release of sequestered calcium ion into cytosol | 3 | 0.837989 | 0.062018 | CD19, GPER1, XCL1 |
| GOTERM_BP_DIRECT | GO:1902600~hydrogen ion transmembrane transport | 4 | 1.117318 | 0.064311 | COX7B, COX7C, UQCRQ, COX6C |
| GOTERM_BP_DIRECT | GO:0010468~regulation of gene expression | 5 | 1.396648 | 0.064483 | CRIP1, G6PC, ZNF683, POU5F1, GDNF |
| GOTERM_BP_DIRECT | GO:0048662~negative regulation of smooth muscle cell proliferation | 3 | 0.837989 | 0.070358 | ANG, PPARG, PPARGC1A |
| GOTERM_BP_DIRECT | GO:0007165~signal transduction | 25 | 6.98324 | 0.071543 | STX2, RHPN1, PPARG, ANXA1, NR4A2, KIR2DS2, ARHGAP19, TNFRSF17, RCVRN, CD70, CD72, ARHGAP24, GDNF, SALL3, CLIC3, ITGB1BP2, OR51B5, MAP3K10, RHEB, SYNGAP1, XCL1, EEF1D, XCL2, SRGAP1, NMU |
| GOTERM_BP_DIRECT | GO:0070836~caveola assembly | 2 | 0.558659 | 0.073106 | CAV1, EMP2 |
| GOTERM_BP_DIRECT | GO:0007224~smoothened signaling pathway | 4 | 1.117318 | 0.085944 | IFT172, HIPK2, MAP3K10, GLI1 |
| GOTERM_BP_DIRECT | GO:0046321~positive regulation of fatty acid oxidation | 2 | 0.558659 | 0.087075 | PPARG, PPARGC1A |
| GOTERM_BP_DIRECT | GO:0001765~membrane raft assembly | 2 | 0.558659 | 0.087075 | S100A10, EMP2 |
| GOTERM_BP_DIRECT | GO:2000272~negative regulation of receptor activity | 2 | 0.558659 | 0.087075 | PCSK9, PPARGC1A |
| GOTERM_BP_DIRECT | GO:0042493~response to drug | 9 | 2.513966 | 0.089647 | PAM, SLC8A1, JUN, PPARG, ANXA1, ADAM17, MDM2, UBE2B, PPARGC1A |
| GOTERM_BP_DIRECT | GO:0001890~placenta development | 3 | 0.837989 | 0.097314 | HTRA1, ANG, PPARG |
| GOTERM_BP_DIRECT | GO:0006641~triglyceride metabolic process | 3 | 0.837989 | 0.097314 | CAV1, G6PC, PCSK9 |
| GOTERM_BP_DIRECT | GO:0010524~positive regulation of calcium ion transport into cytosol | 3 | 0.559701 | 0.038958 | CAV1, P2RX3, CD4 |
| GOTERM_BP_DIRECT | GO:0007267~cell-cell signaling | 13 | 2.425373 | 0.039565 | NOV, LALBA, MLN, NAMPT, NRP1, CXCL5, BHLHA15, CCR1, IL1B, ITGB2, CXCL6, FGF3, FGF4 |
| GOTERM_BP_DIRECT | GO:0048010~vascular endothelial growth factor receptor signaling pathway | 6 | 1.119403 | 0.04242 | VEGFC, NRP1, ROCK1, VEGFA, FOXC2, FOXC1 |
| GOTERM_BP_DIRECT | GO:0007399~nervous system development | 14 | 2.61194 | 0.043738 | CPLX2, MOBP, NRN1, DCTN1, PCDH18, NUMBL, HES1, SLC4A10, DOK4, TPP1, HES4, LSAMP, VEGFA, TMOD2 |
| GOTERM_BP_DIRECT | GO:0060038~cardiac muscle cell proliferation | 3 | 0.559701 | 0.045247 | PRKAR1A, FOXC2, FOXC1 |
| GOTERM_BP_DIRECT | GO:0000122~negative regulation of transcription from RNA polymerase II promoter | 28 | 5.223881 | 0.045389 | CAV1, MTDH, SNCA, CNOT2, NR2E3, FLCN, ZBTB18, TCF7L2, SCGB1A1, KANK2, PROP1, EZR, RARA, SOX17, BRMS1L, KLF5, ZNF280D, MLXIPL, STAT1, CDKN1C, HES1, HDAC5, NCOA2, DR1, VEGFA, FOXC2, CUX2, NFIB |
| GOTERM_BP_DIRECT | GO:0030100~regulation of endocytosis | 4 | 0.746269 | 0.048306 | PACSIN1, PACSIN2, ZFYVE16, RAB5A |
| GOTERM_BP_DIRECT | GO:0001975~response to amphetamine | 4 | 0.746269 | 0.048306 | OXT, DRD4, GRIN1, SLC18A2 |
| GOTERM_BP_DIRECT | GO:0034097~response to cytokine | 5 | 0.932836 | 0.048937 | BCL2, SYNJ1, RARA, STAT1, SCGB1A1 |
| GOTERM_BP_DIRECT | GO:0007032~endosome organization | 4 | 0.746269 | 0.052289 | HOOK1, DNAJC13, SNX10, HOOK3 |
| GOTERM_BP_DIRECT | GO:0031663~lipopolysaccharide-mediated signaling pathway | 4 | 0.746269 | 0.052289 | MTDH, IL1B, TLR4, CD6 |
| GOTERM_BP_DIRECT | GO:0042593~glucose homeostasis | 7 | 1.30597 | 0.052332 | BHLHA15, LEPR, ADRA2A, MLXIPL, ADIPOR1, TCF7L2, PCK1 |
| GOTERM_BP_DIRECT | GO:0046331~lateral inhibition | 2 | 0.373134 | 0.052417 | HES1, DLL1 |
| GOTERM_BP_DIRECT | GO:0038190~VEGF-activated neuropilin signaling pathway | 2 | 0.373134 | 0.052417 | NRP1, VEGFA |
| GOTERM_BP_DIRECT | GO:1902257~negative regulation of apoptotic process involved in outflow tract morphogenesis | 2 | 0.373134 | 0.052417 | FOXC2, FOXC1 |
| GOTERM_BP_DIRECT | GO:1902336~positive regulation of retinal ganglion cell axon guidance | 2 | 0.373134 | 0.052417 | NRP1, VEGFA |
| GOTERM_BP_DIRECT | GO:0090259~regulation of retinal ganglion cell axon guidance | 2 | 0.373134 | 0.052417 | NRP1, VEGFA |
| GOTERM_BP_DIRECT | GO:0097102~endothelial tip cell fate specification | 2 | 0.373134 | 0.052417 | NRP1, DLL1 |
| GOTERM_BP_DIRECT | GO:2001244~positive regulation of intrinsic apoptotic signaling pathway | 4 | 0.746269 | 0.056426 | CAV1, BOK, BCL2, FLCN |
| GOTERM_BP_DIRECT | GO:0045747~positive regulation of Notch signaling pathway | 4 | 0.746269 | 0.056426 | HES1, NOV, DLL1, LFNG |
| GOTERM_BP_DIRECT | GO:0002042~cell migration involved in sprouting angiogenesis | 3 | 0.559701 | 0.058835 | NRP1, VEGFA, NR4A1 |
| GOTERM_BP_DIRECT | GO:0035050~embryonic heart tube development | 3 | 0.559701 | 0.058835 | FOXC2, FOXC1, SOX17 |
| GOTERM_BP_DIRECT | GO:0001503~ossification | 6 | 1.119403 | 0.0616 | ALOX15, BCL2, FOXC2, FOXC1, RUNX1, AHSG |
| GOTERM_BP_DIRECT | GO:0015031~protein transport | 17 | 3.171642 | 0.063273 | PLEKHM1, MYH9, LMAN1, HOOK3, SEC16B, HOOK1, HSP90B1, DNAJC13, RAB5A, RAB11A, RAB6B, EXOC5, AP5M1, SNAP23, SLC15A4, RAB20, SNX10 |
| GOTERM_BP_DIRECT | GO:0048488~synaptic vesicle endocytosis | 3 | 0.559701 | 0.06609 | PACSIN1, SYNJ1, SNCA |
| GOTERM_BP_DIRECT | GO:0010832~negative regulation of myotube differentiation | 3 | 0.559701 | 0.06609 | HDAC5, NOV, BHLHA15 |
| GOTERM_BP_DIRECT | GO:0030322~stabilization of membrane potential | 3 | 0.559701 | 0.06609 | KCNK15, KCNK7, KCNK10 |
| GOTERM_BP_DIRECT | GO:0030097~hemopoiesis | 5 | 0.932836 | 0.071404 | SH2B3, DLL1, RUNX1, FLCN, ADD2 |
| GOTERM_BP_DIRECT | GO:0071407~cellular response to organic cyclic compound | 5 | 0.932836 | 0.071404 | KLF5, ALPL, P2RY13, IL1B, STAT1 |
| GOTERM_BP_DIRECT | GO:0060749~mammary gland alveolus development | 3 | 0.559701 | 0.073623 | PHB2, VEGFA, EGF |
| GOTERM_BP_DIRECT | GO:0030033~microvillus assembly | 3 | 0.559701 | 0.073623 | KLF5, EZR, RAP2C |
| GOTERM_BP_DIRECT | GO:0071310~cellular response to organic substance | 3 | 0.559701 | 0.073623 | BCL2, IL1B, CUX2 |
| GOTERM_BP_DIRECT | GO:0050860~negative regulation of T cell receptor signaling pathway | 3 | 0.559701 | 0.073623 | EZR, CEACAM1, THY1 |
| GOTERM_BP_DIRECT | GO:0030193~regulation of blood coagulation | 3 | 0.559701 | 0.073623 | CAV1, F2RL1, SCARA5 |
| GOTERM_BP_DIRECT | GO:0034765~regulation of ion transmembrane transport | 7 | 1.30597 | 0.075431 | KCNA10, KCND3, KCNK15, KCNK7, KCNH6, CACNA1E, KCNK10 |
| GOTERM_BP_DIRECT | GO:0098903~regulation of membrane repolarization during action potential | 2 | 0.373134 | 0.077588 | CAV1, CACNB3 |
| GOTERM_BP_DIRECT | GO:0019065~receptor-mediated endocytosis of virus by host cell | 2 | 0.373134 | 0.077588 | CAV2, CAV1 |
| GOTERM_BP_DIRECT | GO:0045608~negative regulation of auditory receptor cell differentiation | 2 | 0.373134 | 0.077588 | HES1, DLL1 |
| GOTERM_BP_DIRECT | GO:2000286~receptor internalization involved in canonical Wnt signaling pathway | 2 | 0.373134 | 0.077588 | CAV1, RAB5A |
| GOTERM_BP_DIRECT | GO:1902896~terminal web assembly | 2 | 0.373134 | 0.077588 | EZR, VIL1 |
| GOTERM_BP_DIRECT | GO:2001170~negative regulation of ATP biosynthetic process | 2 | 0.373134 | 0.077588 | PID1, FLCN |
| GOTERM_BP_DIRECT | GO:0061009~common bile duct development | 2 | 0.373134 | 0.077588 | HES1, SOX17 |
| GOTERM_BP_DIRECT | GO:0033280~response to vitamin D | 3 | 0.559701 | 0.081415 | ALPL, PTGS2, CD4 |
| GOTERM_BP_DIRECT | GO:0002690~positive regulation of leukocyte chemotaxis | 3 | 0.559701 | 0.081415 | CXCL5, F2RL1, CXCL6 |
| GOTERM_BP_DIRECT | GO:0048169~regulation of long-term neuronal synaptic plasticity | 3 | 0.559701 | 0.081415 | GRIN1, SNCA, RAB11A |
| GOTERM_BP_DIRECT | GO:0045022~early endosome to late endosome transport | 3 | 0.559701 | 0.081415 | HOOK1, RAB5A, HOOK3 |
| GOTERM_BP_DIRECT | GO:0001756~somitogenesis | 4 | 0.746269 | 0.084296 | FOXC2, DLL1, FOXC1, LFNG |
| GOTERM_BP_DIRECT | GO:0010628~positive regulation of gene expression | 12 | 2.238806 | 0.091402 | SEC16B, PID1, CAV1, EZR, DDX3X, CD46, VEGFA, IL1B, TLR4, EDAR, CUX2, EPB41L4B |
| GOTERM_BP_DIRECT | GO:0060326~cell chemotaxis | 5 | 0.932836 | 0.094163 | NOV, CXCL5, CXCL6, FPR2, DOCK4 |
| GOTERM_BP_DIRECT | GO:0008217~regulation of blood pressure | 5 | 0.932836 | 0.094163 | SGK1, PTGS2, DLL1, NPPA, GCH1 |
| GOTERM_BP_DIRECT | GO:0051384~response to glucocorticoid | 5 | 0.932836 | 0.094163 | ALPL, PTGS2, BCL2, OXT, SCGB1A1 |
| GOTERM_BP_DIRECT | GO:0046330~positive regulation of JNK cascade | 5 | 0.932836 | 0.094163 | TAOK2, F2RL1, IL1B, TLR4, EDAR |
| GOTERM_BP_DIRECT | GO:0070374~positive regulation of ERK1 and ERK2 cascade | 9 | 1.679104 | 0.095352 | ALOX15, NRP1, PHB2, CCR1, VEGFA, F2RL1, CHI3L1, TLR4, FGF4 |
